## Supplementary material for "Timeless couples G quadruplex detection with processing by DDX11 during DNA replication": Combined supplementary information

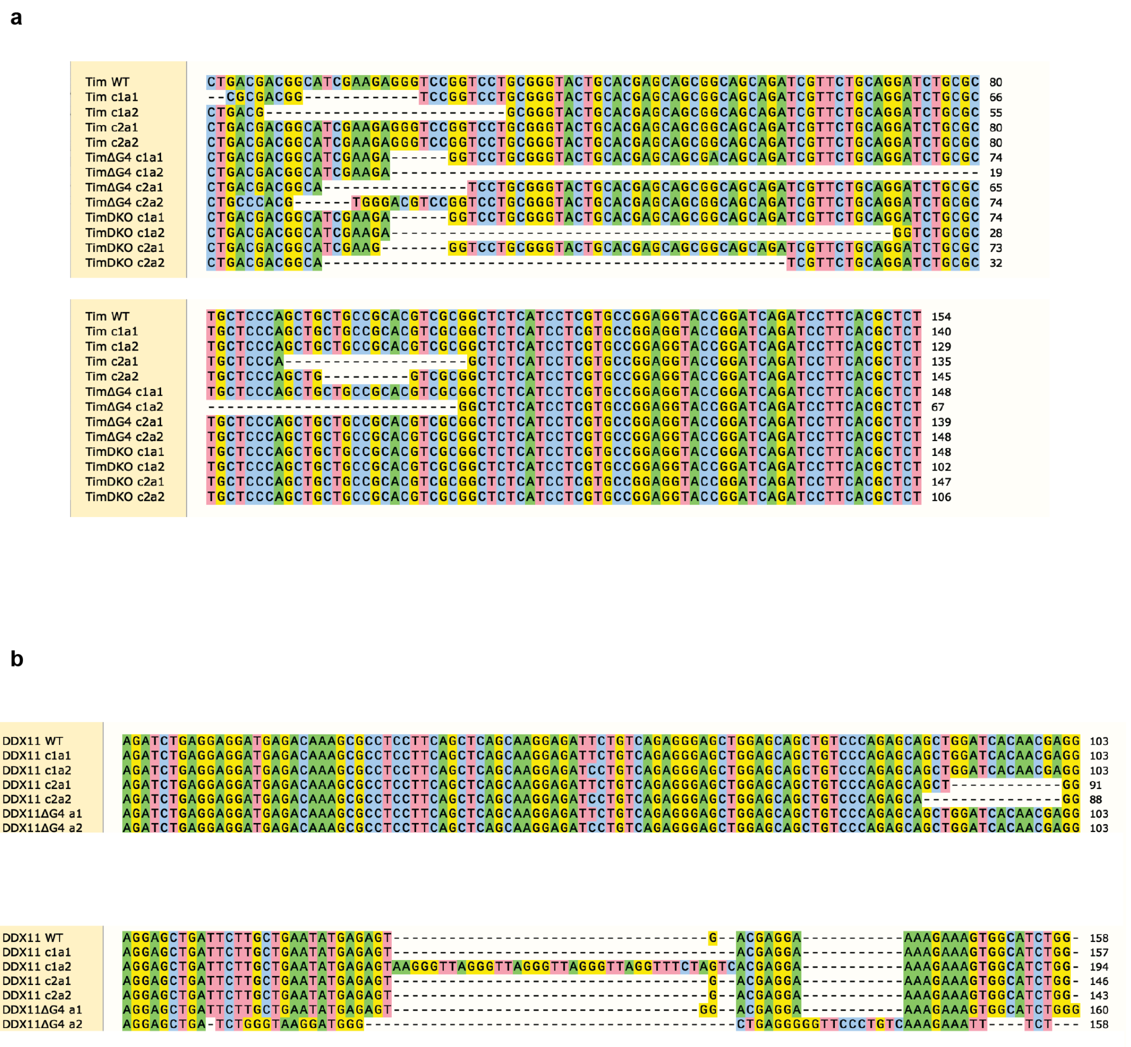


**Supplementary Fig. 1. CRISPR-Cas9 induced deletions in *TIMELESS* and *DDX11.* A.** Aligned sequences of Exon 1 of the chicken *TIMELESS* locus showing the disruptions induced in the different clones used in this study. a1 and a2 refer to the two alleles. Note the telomeric repeat inserted in clone DDX11 c1a2. All disruptions lead to loss of frame.

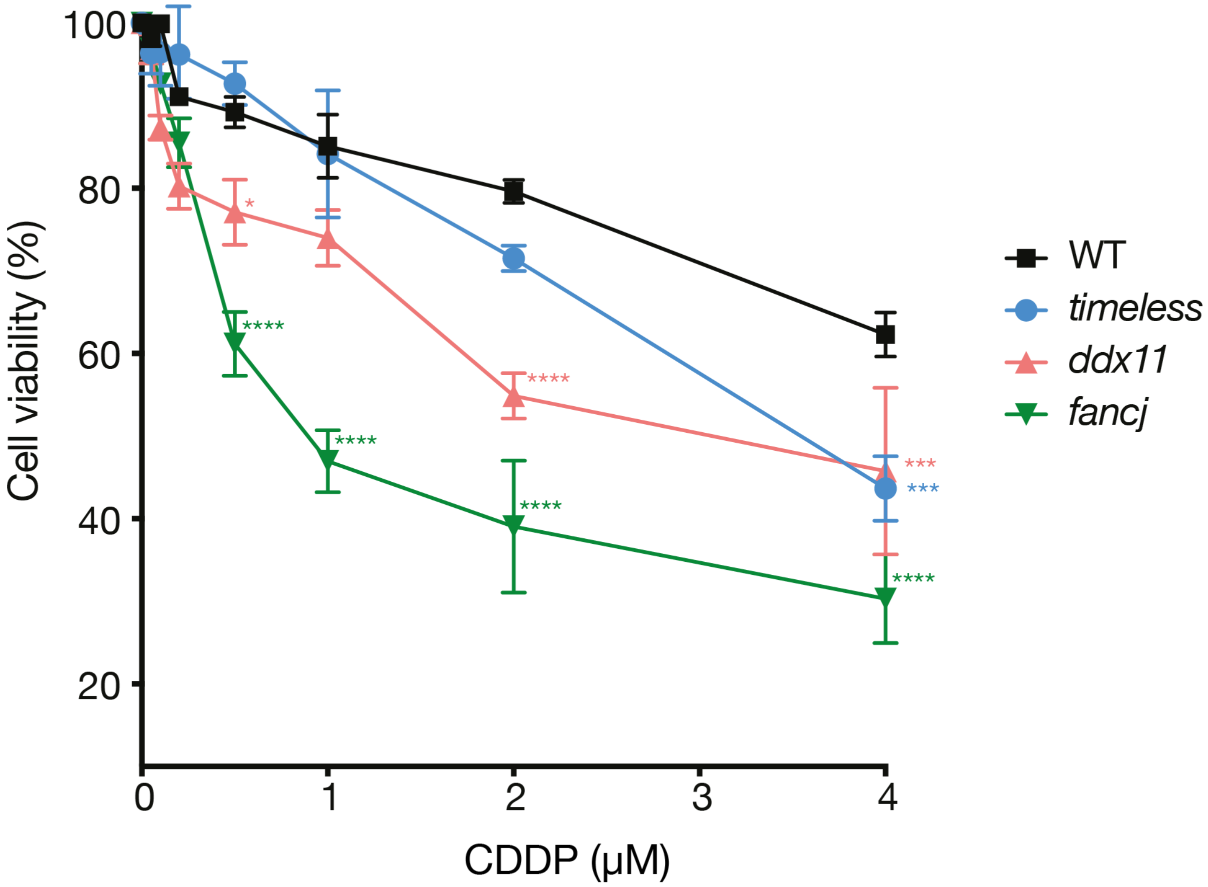


**Supplementary Fig. 2. Sensitivity of wild type (WT), *timeless*, *ddx11* and *fancj* DT40 mutants to cisplatin (CDDP)**. Cell viability, assessed by MTS assay, of DT40 wild type, *ddx11*, *timeless*, *fancj*, after 72 h in presence of cisplatin at the indicated doses. The values represent the means (error bars indicate SD) of two independent experiments performed in triplicate. * $p < 0.05$, *** $p < 0.001$ and **** $p < 0.0001$; one-way ANOVA compared to the wild type.

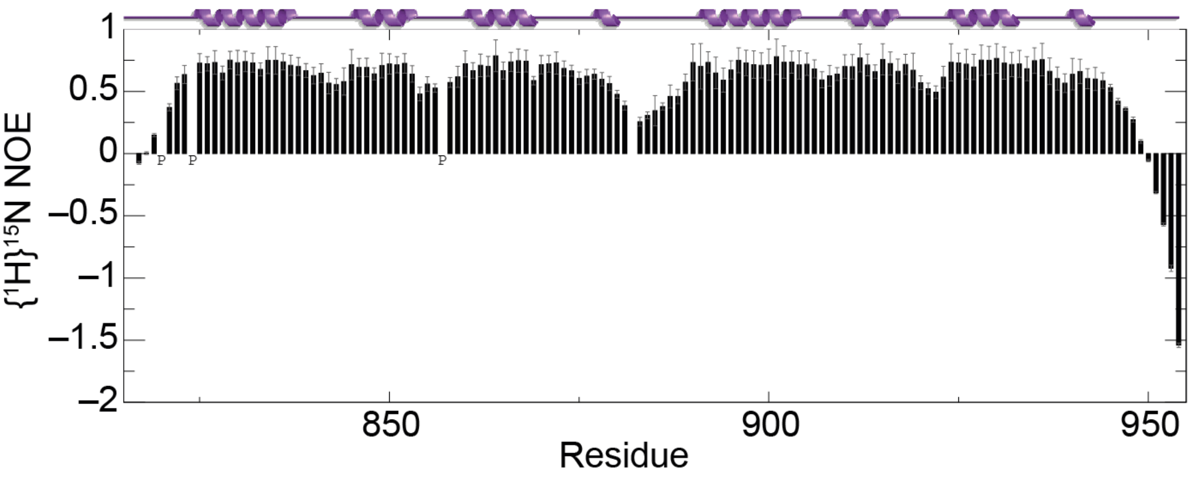


**Supplementary Fig. 3. Backbone dynamics of Timeless 816-954.** $^{1H15}$N heteronuclear nuclear Overhauser enhancement (NOE) values for the inter-domain linker region (residues 881-890) showed evidence of increased motions (NOE < 0.6) on a time scale faster than the overall tumbling rate. The linker is therefore flexible and confers independent mobility on the domains, which leads to a lack of global convergence of the NMR ensemble. Proline residues that lack $^1H^N$ are marked 'P'; the $^1H^N$ of residue 882 was not detected due to conformational exchange broadening. Secondary structure from PDBsum.

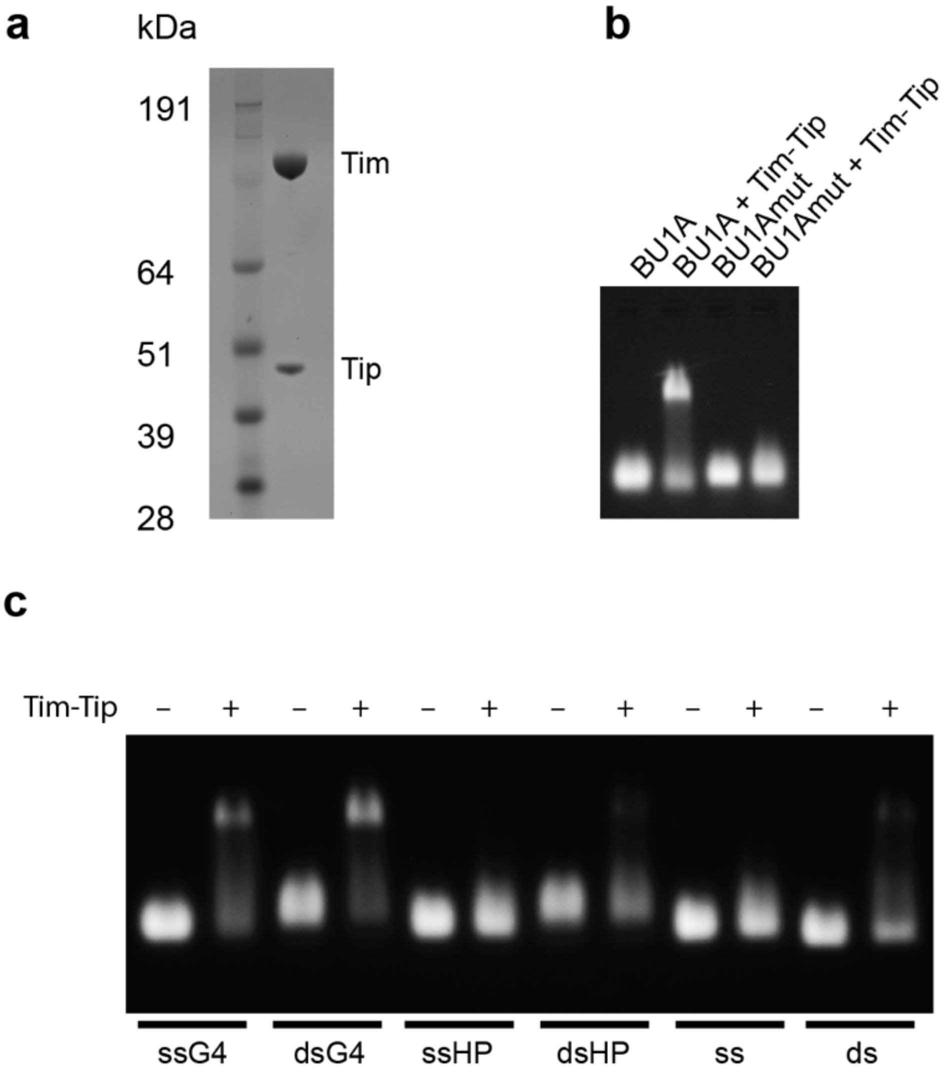


**Supplementary Fig. 4. The Timeless-Tipin complex shows a preference for binding G-quadruplex DNA structures.** A Coomassie-stained SDS-PAGE gel of purified Timeless-Tip complex. B Electrophoretic mobility shift assay (EMSA) showing the binding of Timeless-Tipin to G-quadruplex sequence BU1A+3.5. Mutation of the G-quadruplex sequence (BU1A+3.5mut) disrupts Timeless-Tipin binding (see Supplementary Table 2 for sequence details). Timeless-Tipin and DNA are both present at a final concentration of 5 μM. C EMSA showing the binding of Timeless-Tipin to G-quadruplex sequences (ssG4, dsG4) but not single-stranded DNA (ss), double-stranded DNA (ds) or hairpin-containing sequences (ssHP, dsHP). Timeless-Tipin and DNA are both present at a final concentration of 5 μM.

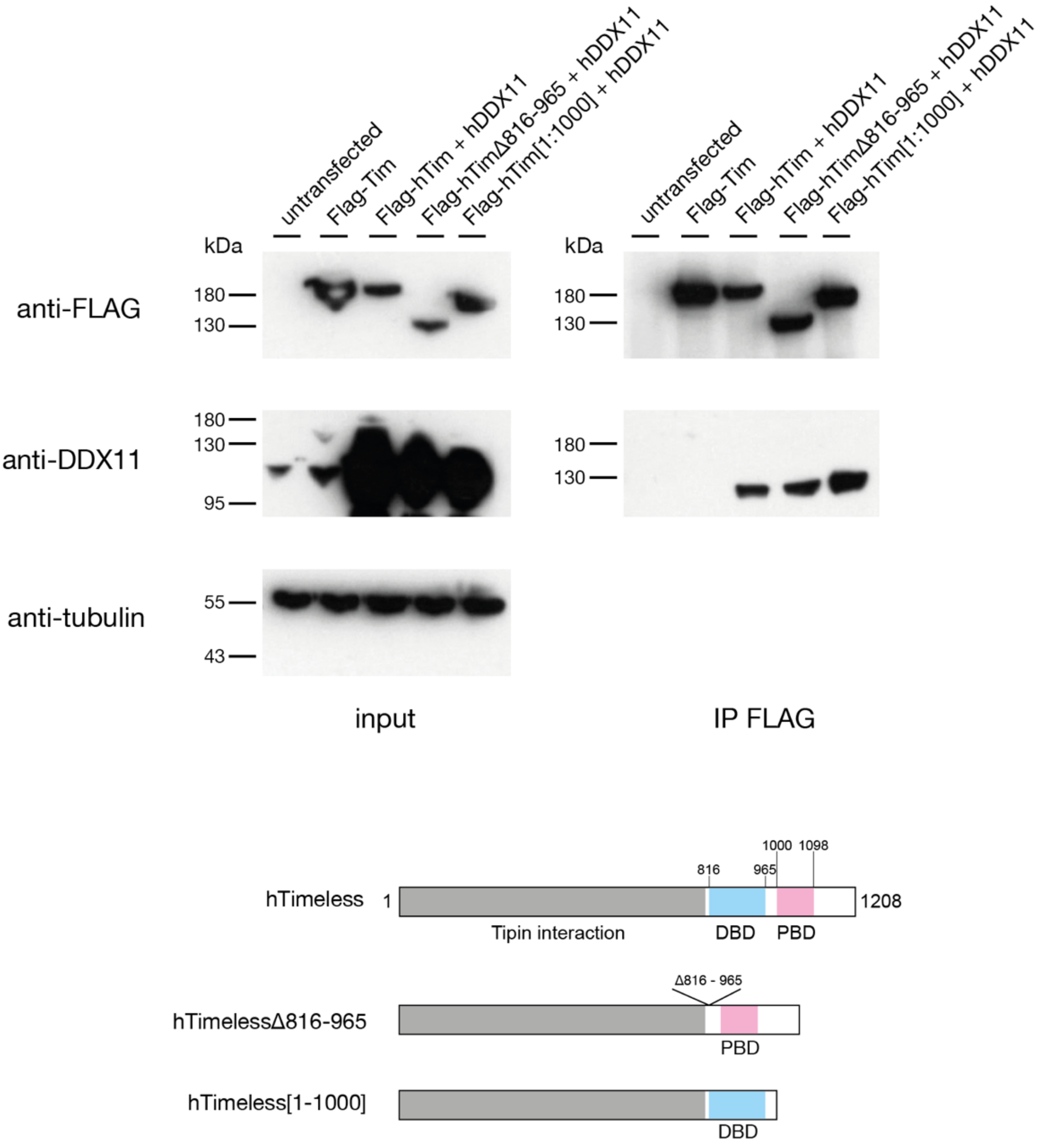


**Supplementary Fig. 5. The C-terminus of Timeless is not required for its interaction with DDX11.** HEK293T cells were transiently transfected with a plasmid encoding Flag-hTimeless, or co-transfected with plasmids encoding hDDX11 and Flag-Timeless or with Timeless mutated to delete the DNA binding domain (ΔDBD: deletion of region S816-S965) or PARP binding domain (PARP*: truncation at V1000). 24 h after transfection, whole cell extracts were subjected to immuno-precipitation with anti-Flag magnetic beads. Western Blot analyses were performed to detect overexpressed DDX11 protein in the pulled down samples using a specific antibody. Upper panel: Input and pulled down samples transfected with different Timeless constructs detected with an anti-Flag antibody. Bottom panel: Input and pulled down samples transfected with different Timeless constructs detected with an anti-DDX11 antibody. Tubulin was used as a loading control for the input samples.

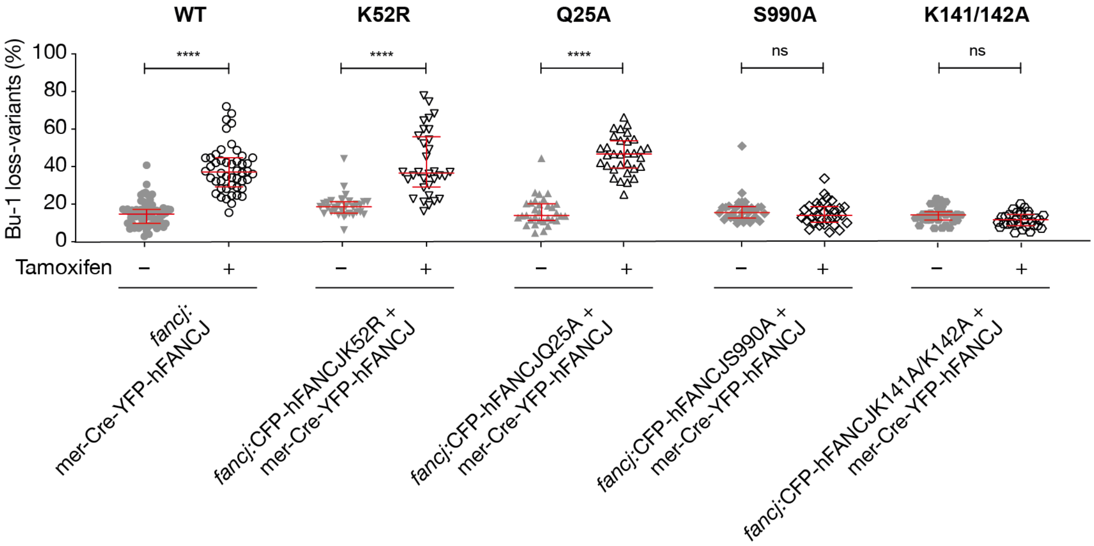


**Supplementary Fig. 6. The catalytic activity of FANCJ is required for its role in suppressing G4-induced instability of *BU-1* expression.** Fluctuation analysis for the generation of Bu-1 loss variants in an inducible system to study FANCJ function (see Methods for full details). Briefly FANCJ-deficient DT40 cells are rescued with two transgenes, one encoding the wild type protein, the other the mutant in question. The wild type transgene is flanked by loxP sites and can be deleted by expression of Cre recombinase, induced by treatment of cells with tamoxifen. The K52R and Q25A mutants of FANCJ both disrupt the helicase activity of the enzyme [1,2]. S990A disrupts the interaction of FANCJ with BRCA1, which is important for the role of FANCJ in homologous recombination [3]. K141/142A disrupts the interaction of FANCJ with MutLα, which is needed for efficient interstrand crosslink repair [4].

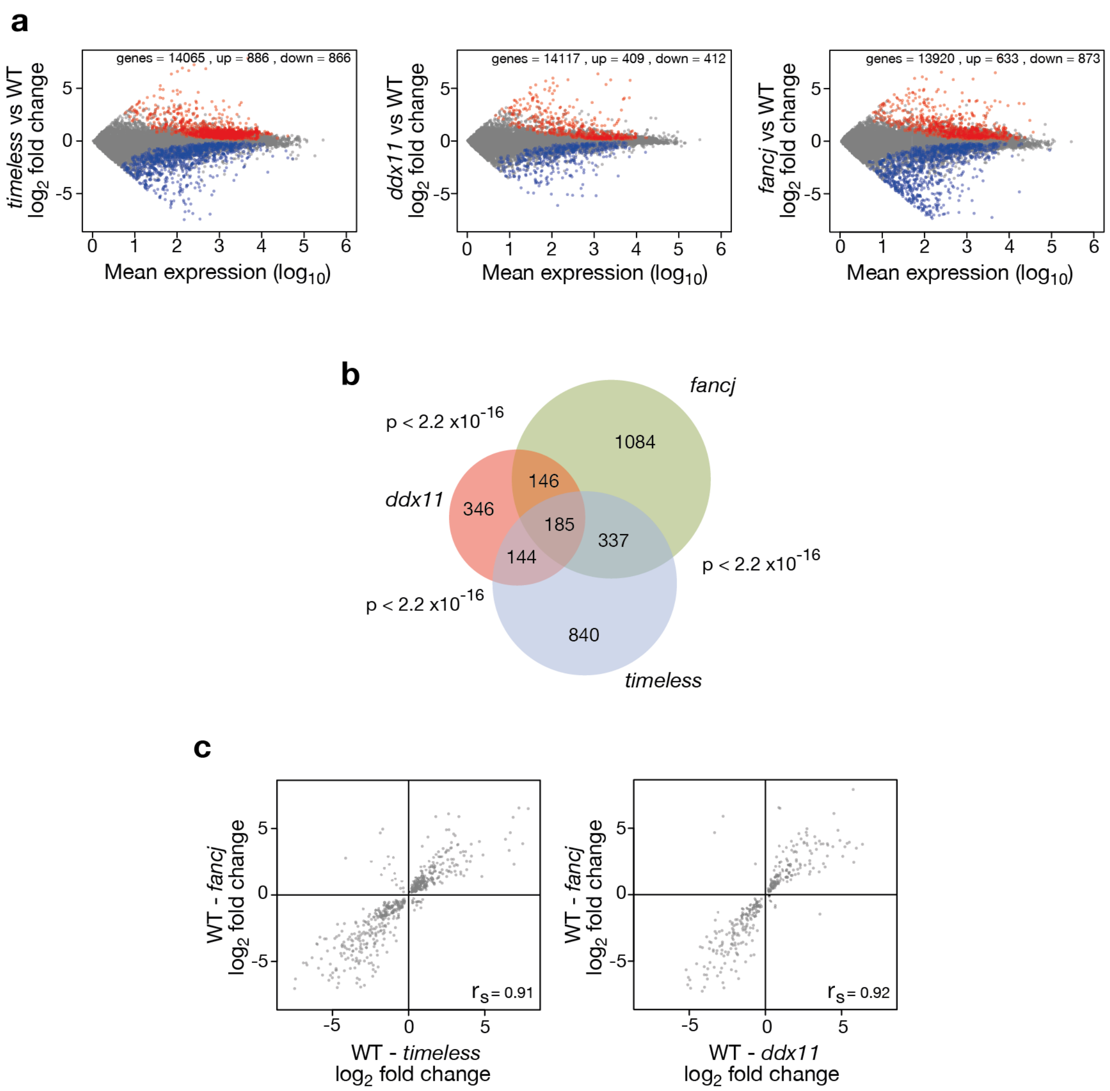


**Supplementary Fig. 7. Gene expression dysregulation in *timeless* and *ddx11* DT40 cells.** **a.** Dysregulated genes in *timeless* (left panel), *ddx11* (centre panel) and *fancj* (right panel) mutants relative to wild type. All genes with > 1 transcript per million in both conditions are plotted. Significantly ($p \geq 0.95$) upregulated genes shown in red; downregulated in blue **b.** Venn diagram showing the overlap in genes deregulated in *timeless*, *ddx11* and *fancj* relative to wild type. $p < 2.2 \times 10^{-16}$ for each pairwise comparison (Fisher hypergeometric distribution). **c.** Correlation of magnitude and direction of change of genes dysregulated (relative to wild type) in *fancj vs. timeless* (left panel) and *dd*x11 (right panel) DT40 cells. $r_s$ (Spearman rho) is shown for each correlation.

**Supplementary Table 1.** Oligos used for molecular cloning, site-directed mutagenesis and CRISPR/Cas9 gene disruption

| Identifier | 5′-3′ DNA Sequence |
|---|---|
| hTimΔDBD-D965-ClaI-GA-Fw | AGATGACTGAGGGCTATGGCTCCCTGGATGACAGGTCTTCCATCGATTTTTGCCAGGAAGATCTGGAAGAAGAGG |
| hTimΔDBD-S816-ClaI-GA-Rev | TTCCTCAGGCAGGTTTTCCTCTTCTTCCAGATCTTCCTGGCAAAAATCGATGGAAGACCTGTCATCCAGGGAGCC |
| hTimPBDtrunc-pcDNA3.1-bGH-TGA-NotI-GA-Fw | CAAGTCCAGGGTAGCTTAGTCTGAGCGGCCGCTCGAGTCTAGAGGGCCCTTCG |
| hTimPBDtrunc-NotI-GA Rev | CCCTCTAGACTCGAGCGGCCGCTCAGACTAAGCTACCCTGGACTTGTTCTGC |
| hTimC-tertrunc-pcDNA3.1-bGH-TGA-NotI-GA-Fw | GCTCCCTGGATGACAGGTCTTCCTGAGCGGCCGCTCGAGTCTAGAGGGCCCTTCG |
| hTimC-tertrunc-S816-NotI-GA-Rev | AAGGGCCCTCTAGACTCGAGCGGCCGCTCAGGAAGACCTGTTCATCCAGGGAGCC |
| hFANCJ K52R A155G Fw | AAGTAAGGCTAAGCTTCTTCCACTTCCTGTGGGAC |
| hFANCJ K52R A155G Rev | GTCCCACAGGAAGTGGAAGAAGCTTAGCCTTACTT |
| cDDX11 CRISPR gRNA top | CACCGTGCTGAATATGAGAGTGACG |
| cDDX11 CRISPR gRNA bottom | AAACCGTCACTCTCATATTCAGCAC |
| cTim CRISPR (KO) gRNA top | CACCGCGACGGCATCGAAGAGGGTC |
| cTim CRISPR (KO) gRNA bottom | AAACGACCCTCTTCGATGCCGTCGC |

| | |
|---|---|
| cTim CRISPR (C-ter truncation) gRNA top | CACCGACTTGTCGTCGTGTGCGGCC |
| cTim CRISPR (C-ter truncation) gRNA top | AAACGGCCGCACACGACGACAAGTC |

**Supplementary Table 2.** DNA sequences used in Timeless DNA-binding experiments

| Identifier | 5′-3′ DNA sequence (secondary structure forming elements are highlighted in red) |
|---|---|
| ssGQ | **6FAM-** ACGAGAGCTAGCACATTTTGA**GGG**T**GGG**TA**GGG**T**GGG**TAATTTTCACGTAGAACCTGT |
| dsGQ | ssGQ+ ACAGGTTCTACGTGATTTTTTTTTTTTTTTTTTTTTTTTTTTTTGTGCTAGCTCTCGT |
| ssHP | **6FAM-** ACGAGAGCTAGCACATT**TTGAGGCTGCG**TTT**CGCAGCCTCAA**TTTCACGTAGAACCTGT |
| dsHP | ssHP+ ACAGGTTCTACGTGATTTTTTTTTTTTTTTTTTTTTTTTTTTTTGTGCTAGCTCTCGT |
| ss | **6FAM-** ACGAGAGCTAGCACATTTTGAGTGTCAGTAGCGTCTGTAATTTTCACGTAGAACCTGT |
| ds | ss+ ACAGGTTCTACGTGAAAATTACAGACGCTACTGACACTCAAAATGTGCTAGCTCTCGT |
| BU1 +3.5 [5] | **6FAM-** AGCTAGCACATTTTAA**GGG**CT**GGG**T**GGG**TGCTGTCAA**GGG**CT**GGG**TTTTCACGTAGAA |
| BU1 +3.5mut [5] | **6FAM-** AGCTAGCACATTTTAAGTTCTGTTTGTTTGCTGTCAAGTTCTGTTTTTTCACGTAGAA |
| 2JPZ [6] | **6FAM**-TTA**GGG**TTA**GGG**TTA**GGG**TTA**GGG**TT |
| 1XAV [7] | **6FAM**-TGA**GGG**T**GGG**TA**GGG**T**GGG**TAA |
| G4#2 [5] | **6FAM**-TAAT**GGG**TTT**GGG**TTT**GGG**TTT**GGG**T |
| G4#4 [5] | **6FAM**-TAATTTT**GGG**T**GGG**T**GGG**T**GGG**TTTT |
| 2O3M [8] | **6FAM**-TAA**GGG**A**GGG**CGCT**GGG**AGGA**GGG** |
| Bcl2Mid [9] | **6FAM**-ATA**GGG**CGC**GGG**AGGAGG**GGG**C**GGG** |
| ρ-globin [10] | **6FAM**-TAA**GGGG**AGTAAAA**GGG**AGC**GGGG**TGCT**GGG** |

**Supplementary Table 3. Data collection and refinement statistics for Tim DBD C-term (residues 883-947).**

| | |
|---|---|
| *Data collection* | |
| Wavelength | 0.9686 Å |
| Resolution range (Å) | 31.46 - 1.15 (1.191 - 1.15) |
| Space group | $P6_5$ |
| Unit cell | 55.407 55.407 41.659 90 90 120 |
| Total reflections | 104424 (3064) |
| Unique reflections | 25072 (1920) |
| Multiplicity | 4.2 (1.6) |
| Completeness (%) | 96.33 (74.84) |
| Mean I/sigma(I) | 6.82 (1.36) |
| Wilson B-factor | 11.30 |
| R-merge | 0.1377 (0.4873) |
| R-meas | 0.1543 (0.6505) |
| R-pim | 0.06851 (0.4274) |
| CC1/2 | 0.975 (0.464) |
| CC* | 0.994 (0.796) |
| *Refinement* | |
| Reflections used in refinement | 24985 (1922) |
| Reflections used for R-free | 1267 (101) |
| R-work | 0.1280 (0.2681) |
| R-free | 0.1548 (0.3632) |
| CC(work) | 0.973 (0.809) |

| | |
|---|---|
| CC(free) | 0.963 (0.769) |
| Number of non-hydrogen atoms | 659 |
| macromolecules | 527 |
| solvent | 132 |
| Protein residues | 62 |
| RMS(bonds) | 0.008 |
| RMS(angles) | 0.96 |
| Ramachandran favored (%) | 96.67 |
| Ramachandran allowed (%) | 3.33 |
| Ramachandran outliers (%) | 0.00 |
| Rotamer outliers (%) | 0.00 |
| Clashscore | 1.86 |
| Average B-factor | 17.72 |
| macromolecules | 14.14 |
| solvent | 32.01 |

Statistics for the highest-resolution shell are shown in parentheses.

**Supplementary Table 4.** Summary of the restraints used in the calculation of the Timeless 816-954 structure and characterisation of the ensemble of energy-minimised structures.

| | | |
|---|---|---|
| *NOE upper distance limits* | | |
| Total | 4730 | |
| Ambiguous | 1912 | |
| Unambiguous | 2818 | |
| *Dihedral angle constraints* | | |
| Total | 278 | |
| *Residual NOE violations* (Å) | | |
| Number ≥ 0.5 | 0 | |
| Number ≥ 0.1 | 117 | |
| *Residual angle violations* (deg.) | | |
| Number ≥ 2.0 | 0 | |
| *Energies* (kcal $mol^{-1}$) | | |
| Total | –4691 ± 40 | |
| van der Waals | –1348 ± 40 | |
| Electrostatic | –5207 ± 53 | |
| *Ramachandran statistics (% residues)* | | |
| Core regions | 83.7 | |
| Allowed regions | 16.3 | |
| Generously allowed regions | 0 | |
| Disallowed regions | 0 | |
| *r.m.s.d. from ideal geometry* | | |
| Bond lengths (Å) | 0.0052 ± 0.0002 | |
| Bond angles (deg.) | 0.660 ± 0.008 | |
| *r.m.s.d. to mean coordinates* (Å) | *Backbone* | *Heavy atoms* |
| Protein (824-880) | 0.494 | 0.580 |
| Protein (891-944) | 0.487 | 0.527 |

For the analysis, 20 water-refined, energy-minimised structural conformations were used. Ramachandran statistics were calculated using PDBsum.
